# A comparison of deep learning and linear-nonlinear cascade approaches to neural encoding

**DOI:** 10.1101/463422

**Authors:** Theodore H. Moskovitz, Nicholas A. Roy, Jonathan W. Pillow

## Abstract

A large body of work on neural encoding has focused on “cascade” type models such as the linear-nonlinear-Poisson (LNP) model. This approach seeks to describe the encoding process in terms of a series of stages: (1) projection of the stimulus onto a bank of linear filters; (2) a nonlinear function combining these filter outputs; and (3) a noisy spike generation process. Here we explore the relationship of the LNP modeling framework to more recent approaches arising from the deep learning literature. Specifically, we show that deep neural network (DNN) and convolutional neural network (CNN) models of neural activity sit firmly within the LNP framework, and correspond to particular parametrizations of the nonlinear stage of the LNP model. Using data from primate retina and primary visual cortex, we compare the performance of LNP models fit with deep learning methods to LNP models fit with traditional estimators, including spike-triggered covariance (STC), information-theoretic spike-triggered averaging and covariance (iSTAC), and maximum likelihood estimators also known as “maximally informative dimensions” (MID). We show that models with nonlinearities parametrized by deep networks achieve higher accuracy for a fixed number of filters, and can extract a larger number of informative filters than traditional models. Finally, we perform a dimensionality analysis of LNP models trained with deep learning methods, revealing that a large number of filters are needed to accurately describe the neural responses of many cells even early in the visual pathway. This result overturns one of the central tenets of the LNP modeling framework: that neural computations are low-dimensional, or depend on the stimulus only via its projection onto a small number of linear filters. We discuss the implications of these findings for both the fitting and interpretation of LNP encoding models.

## 1 Introduction

One of the fundamental problems in computational neuroscience is characterizing how neurons convert sensory inputs to spike responses. This is commonly referred to as the *neural encoding problem*. The difficulty of this problem arises from the stochasticity of neural responses as well as the high dimensionality of sensory stimuli, which makes it impossible to explore the entire space of possible inputs to a neuron (e.g., the space of all 2D images). An important simplifying assumption that has made the neural coding problem tractable is the idea that neurons compute their response in a low-dimensional space, i.e., that their responses depend on a low-dimensional projection of the stimulus (de Ruyter van Steveninck & Bialek, 1988; Bialek, Rieke, de Ruyter van Steveninck, & Warland, 1991; Schwartz, Chichilnisky, & Simoncelli, 2002; Aguera y Arcas & Fairhall, 2003; Aguera y Arcas, Fairhall, & Bialek, 2003; Sharpee, Rust, & Bialek, 2004; Rust, Schwartz, Movshon, & Simoncelli, 2005; Schwartz, Pillow, Rust, & Simoncelli, 2006; Aljadeff, Lansdell, Fairhall, & Kleinfeld, 2016). Most approaches to neural characterization therefore focus primarily on identifying the subspace of the sensory stimulus space that affects a neuron’s activity. For example, information-theoretic estimators explicitly seek to identify a small set of input filters that maximize the response-relevant information contained in the stimulus. Similarly, moment-based estimators, such as those using the spike-triggered average (STA) and covariance (STC), directly compute filters that span the principal axes of the spike-triggered stimulus distribution.

These methods come with their own sets of advantages and disadvantages. Moment-based estimators generally have low computational cost but succeed only for restricted stimulus domains and modeling assumptions, while the reverse is true for maximum likelihood or information-based estimators (Williamson, Sahani, & Pillow, 2015). While a large literature has focused on the estimation of linear-nonlinear-Poisson (LNP) encoding models (Korenberg & Hunter, 1986; Sharpee et al., 2004; Paninski, 2004; Truccolo, Eden, Fellows, Donoghue, & Brown, 2005; Schwartz et al., 2006; Gerwinn, Macke, & Bethge, 2010; Park & Pillow, 2011; Williamson et al., 2015; Heitman et al., 2016; Aljadeff et al., 2016), recent work has focused on efforts to model neural responses using deep neural networks (DNNs) (Batty et al., 2017; McIntosh, Maheswaranathan, Nayebi, Ganguli, & Baccus, 2016). Here we show that these two approaches need not be considered distinct, as a DNN with Poisson output noise represents a particular parametrization of an LNP model.

Our paper makes the following contributions: (i) We clarify the theoretical equivalence of the LNP and DNN frameworks. (ii) We investigate differences between the filters obtained by traditional LNP model estimators and DNN-based estimators; (iii) We show that DNN-based LNP models outperform LNP models estimated with traditional estimators, even for simple white noise stimuli; (iv) We show that early visual neural responses are in fact high-dimensional, in contrast to previous assumptions, and that DNN-based LNP models are better able to take advantage of this fact.

## 2 Background: Linear-Nonlinear-Poisson model

The linear-nonlinear-Poisson (LNP) model describes the neural encoding process in terms of a series of three stages (Schwartz et al., 2006; Pillow, 2007). First, a linear stage projects the high-dimensional sensory stimulus onto a set of linear filters, performing dimensionality reduction to a neural feature space in which the neuron is assumed to compute its response. Second, a nonlinear stage maps the filter outputs to a non-negative spike rate. Third, the spike rate drives spiking via an inhomogeneous Poisson process, which is typically sampled in discrete time bins. (See Figure 1 for an LNP model schematic).

**Figure 1:**
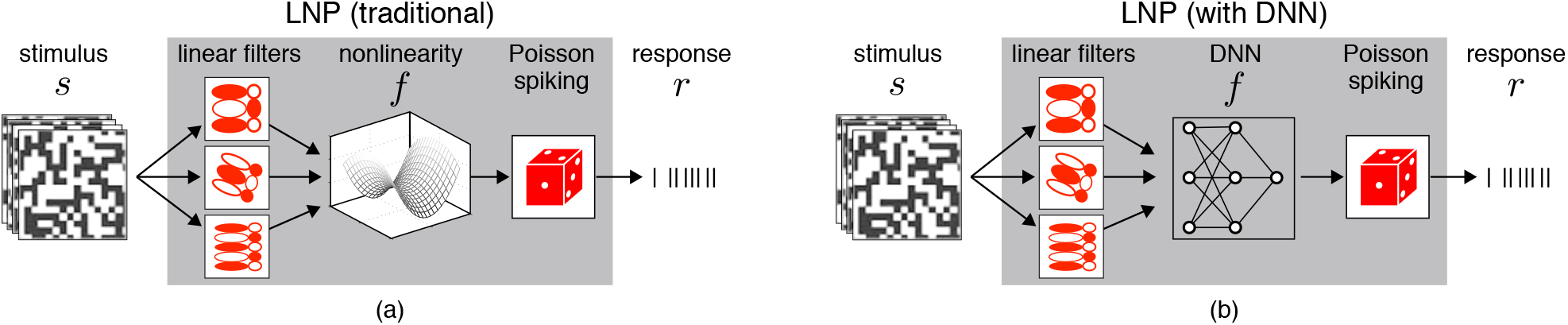
Neural encoding frameworks. (a) The classic multi-filter LNP framework, in which a small set of linear filters *K* perform dimensionality reduction on the input stimulus. The set of filter responses in feature space is then passed to a nonlinear function *f* whose output *λ* is interpreted as a Poisson spiking rate. (b) A DNN model for neural encoding works equivalently, with the input filtered through a linear transformation before being passed to a multilayer nonlinear function in the form of a neural network.

The model is parametrized by a set of filters contained in the columns of a matrix, *K* = [**k**_1_,…, **k**_*d*_], and a point-wise nonlinearity *f* : ℝ^*d*^→ ℝ that transforms the *d*-dimensional vector of filter outputs to a Poisson spike rate. The filters operate on an *n*-dimensional stimulus s (e.g., an image with *n* pixels) and the model output is a non-negative integer spike count *r* in each time bin. Mathematically, the model can be written:

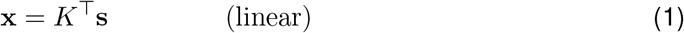

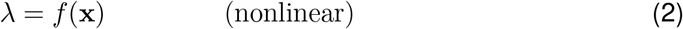

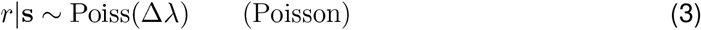

where x is a low-dimensional feature vector resulting from projection of the high-dimensional stimulus s onto the filters in *K*, λ is the instantaneous spike rate, Δ is the time bin width, and the number of spikes *r* follows a Poisson distribution:

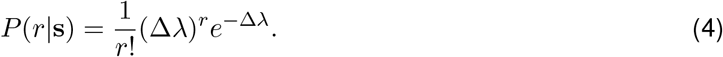

Estimators for the LNP model can be categorized into three classes based on the kind of optimization problem required to obtain the filter estimates:

1. closed-form expressions using sufficient statistics
2. numerical optimization using sufficient statistics
3. numerical optimization requiring multiple passes over the full dataset.

Class 1 includes moment-based estimators like the spike-triggered average (deBoer & Kuyper, 1968; Chichilnisky, 2001) and spike-triggered covariance (STC) analysis (de Ruyter van Steveninck & Bialek, 1988; Schwartz et al., 2002), which provide analytic expressions for filter estimates using first and second moments of the stimulus and spike-triggered stimulus distribution. Class 2 involves estimators that require numerical optimization of a nonlinear objective function that can be evaluated using the stimulus and spike-triggered moments. This class includes information-theoretic Spike-Triggered Average and Covariance (iSTAC) Bayesian spike-triggered covariance, and moment-based estimators for generalized quadratic models (GQMs) (Park & Pillow, 2011; Ramirez & Paninski, 2013; Park, Archer, Priebe, & Pillow, 2013). Class 1 and 2 estimators both require the use of stimuli obeying certain regularity conditions (e.g. elliptical symmetry or Gaussianity), and achieve optimality only under certain assumptions about the nonlinearity (e.g. exponential or exponentiated quadratic) (Paninski, 2003a; Park et al., 2013). The payoff for making these assumptions is that estimates require only a single pass through the data to compute moments.

By contrast, LNP model estimators in Class 3 require a full pass through the data for each evaluation of the obective or loss function. The resulting estimators are consistent for arbitrary choices of stimulus; however, the optimization problem is typically non-convex, meaning that it may be difficult to find a global optimum, and is computationally expensive due to the need to take multiple passes through the entire dataset. Estimators in this class include *maximally informative dimensions (MID)* (Sharpee et al., 2004), which is equivalent to maximum likelihood (ML) estimation of the filters under a non-parametric model of the nonlinearity (Williamson et al., 2015). Class 3 also includes generalized linear models (GLMs) (Truccolo et al., 2005), which typically have a convex loss function but require specifying a nonlinearity *a priori* and are restricted to a single linear filter.

In this paper, we will examine classical LNP model estimators from the three classes defined above, and compare them to an estimator based on deep learning.

### 2.1 Closed-form moment-based estimators

Two classic estimators for the filters in an LNP model can be obtained directly from the first and second moment of the spike-triggered stimulus distribution: the spike-triggered average (STA) and the spike-triggered covariance (STC), respectively. The STA is defined as the average fixed-length stimulus preceding a spike:

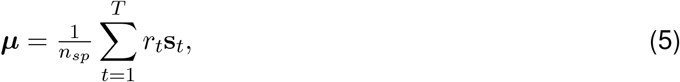

where 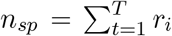 is the total number of spikes. The STA provides an unbiased, consistent estimate of the filter for a single-filter LNP model, assuming that the stimulus distribution (shifted to have zero mean) is spherically symmetric, and the model nonlinearity shifts the mean of the spike-triggered stimulus distribution away from zero (Chichilnisky, 2001; Paninski, 2003b). However, the STA is asymptotically optimal (meaning that it achieves the lowest possible mean-squared error) only in the case that the nonlinearity is exponential, *f* (*x*) = exp(*x* + *a*) (Paninski, 2003b; Pillow & Simoncelli, 2006; Pillow, 2007).

For LNP models with multiple filters, the STA recovers only a single dimension of the subspace spanned by the model filters; additional dimensions can be obtained from eigenvectors of the spike-triggered covariance matrix, defined as

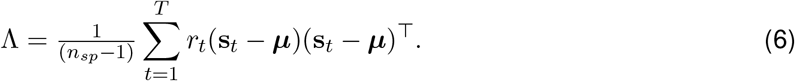

If the stimulus distribution has been whitened so as to have identity covariance, eigenvectors of Λ with eigenvalues significantly different from 1 provide consistent estimators for the filter subspace, provided the stimulus is Gaussian and the nonlinearity shifts the variance of the spike-triggered stimulus distribution along all filter dimensions. (When the stimuli are correlated, the spike-triggered covariance must be adjusted to account for the stimulus covariance matrix; see (Schwartz et al., 2006) for details). The STC-based estimator is computationally inexpensive because it requires only a single pass through the data and an eigendecomposition of the STC matrix. However, it is asymptotically optimal only when the nonlinearity takes the form of an exponentiated quadratic form, *f*(x) = exp(x^⊤^*C*x + *a*) for some matrix *C* and a constant *a* (Pillow & Simoncelli, 2006; Pillow, 2007; Park & Pillow, 2011).

### 2.2 The iSTAC estimator

A second class of LNP model filter estimators rely on optimization of a nonlinear objective function of the spike-triggered moments (STA and STC defined) above. The first such estimator was the information-theoretic Spike-Triggered Average and Covariance (iSTAC) estimator (Pillow & Simoncelli, 2006), which was motivated by the observation that STA and STC-based estimators do not optimally combine information from the first two moments when estimating a filter subspace. For example, for a single-filter model in which the nonlinearity increases both the mean and variance of the spike-triggered stimulus distribution, the filter can be estimated using the STA or the top eigenvector of the STC matrix. However, neither estimate would be optimal because it does not combine information from both moments. The iSTAC estimator overcomes this shortcoming, but at the cost of requiring numerical optimization of a nonlinear objective function. Technically, the objective is the Kullback-Leibler (KL) divergence between Gaussian approximations to the spike-triggered and raw stimulus distributions, which corresponds to an information-theoretic quantity known as the single-spike information (Brenner, Strong, Koberle, Bialek, & de Ruyter van Steveninck, 2000; Sharpee et al., 2004). This objective takes a simple form:

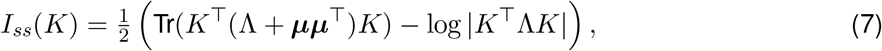

where *I_ss_*(*K*) denotes the single-spike information captured by a filter matrix *K*, and is equal to the KL divergence between *P*(*K*^⊤^x|spike) and *P*(*K*^⊤^x), the spike-triggered and raw stimulus distributions projected onto the filter subspace defined by *K*, respectively. The most informative subspace corresponds to the *K* that maximizes *I_ss_*(*K*) (eq. 7).

The conditions of validity for the iSTAC estimator are similar to those of STC: it is consistent only when the stimulus distribution is Gaussian. However, it achieves asymptotic optimality under slightly more general conditions than STA or STC, namely when the nonlinearity contains exponentiated quadratic and linear forms: *f* (x) = exp(x^*T*^*C*x + b^⊤^x + *a*), for any *C*, b, and *a*. This is an example of a general family of models known as *generalized quadratic models* (GQMs) (Park et al., 2013). Subsequent work has shown that the iSTAC estimator is asymptotically equivalent to a maximum likelihood estimator (Park & Pillow, 2011), and derived generalizations to a larger family of stimulus distributions (Park et al., 2013). One additional advantage of iSTAC over STA and STC-based based estimators is that it comes with closed-form estimators for the parameters of the GQM nonlinearity once the filter matrix estimate is obtained (Pillow & Simoncelli, 2006; Park & Pillow, 2011).

### 2.3 Maximum-likelihood / Infomax estimators

Two important limitations of the estimators defined above is their reliance on spherically symmetric or Gaussian stimuli, and restrictions on the class of nonlinearities for which they achieve optimality. A more powerful if more computationally demanding approach is therefore to perform to joint maximum likelihood inference for the filters *K* and a set of parameters defining the nonlinearity.

An early example of such an approach is the *maximally informative dimensions* (MID) estimator, which used histograms of the projected raw and spike-triggered stimuli to estimate the nonlinearity (Sharpee et al., 2004). Although framed in terms of maximizing single-spike information, later work showed MID to be the maximum likelihood estimator for an LNP model in which the nonlinearity is parametrized by piecewise-constant basis functions (Williamson et al., 2015). In this framework, the nonlinearity can be described as:

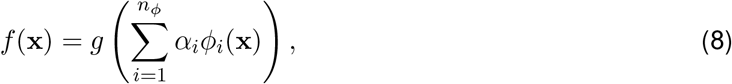

where *ϕ_i_*(·) is the *i*’th basis function (a function that takes a value of 1 within a rectangular region of the filter output space and 0 elsewhere), *α_i_* is the coefficient weighting this basis function, *n_ϕ_* is the number of basis functions, and *g* is a fixed rectifying function (e.g., rectified-linear) that enforces non-negative firing rates. MID corresponds to a joint maximum likelihood estimate for the filter parameters *K* and nonlinearity parameters *α* = (*α*_1_,…, *α_n_ϕ__*), namely:

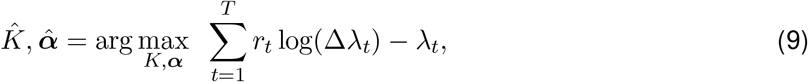

where the right-hand-side is the Poisson log-likelihood (from the log of eq. (4), ignoring constant terms), and *λ_t_* = *f* (*K*^⊤^s_*t*_) is the firing rate computed by the LN cascade applied to stimulus s_*t*_.

Subsequent work proposed using the same maximum likelihood framework with radial or cylindrical basis functions in place of piece-wise constant (“histogram”) basis functions (Williamson et al., 2015). Cylindrical basis functions (CBFs) resemble radial basis functions but are constrained only in some dimensions. A “first-order” CBF, which is constrained in only 1 dimension, is defined

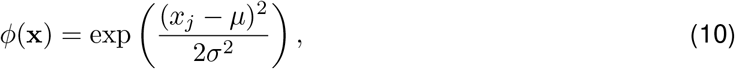

where *x_j_* is a single component of the vector x, and *μ* and *σ* denote the mean and width of the basis function along this dimension. Compared to piece-wise constant basis functions, RBFs and CBFs have the advantage that the nonlinearity is continuous and differentiable, resulting in a log-likelihood function that is continuous and differentiable with respect to the model parameters.

The family of maximum-likelihood estimators defined by equations (8–9) can be considered a *semi-parametric* because they include a parametric component (the filter matrix *K*) and a non-parametric component (a nonlinearity that can be made arbitrarily flexible by growing the set of basis functions as dataset size increases). Statistically, these estimators can achieve asymptotic optimality for any stimulus distribution and neural nonlinearity given appropriate control of the degrees of freedom governing the non-parametric nonlinearity (i.e., number of basis functions). However, in practice these estimators are computationally expensive, due to the need to pass through the entire dataset for each evaluation of the log-likelihood, and may be difficult to fit due to the ubiquity of sub-optimal local optima in the log-likelihood function. These shortcomings provide strong motivation for deep-learning based approaches, which have developed strategies for overcoming precisely these challenges.

### 2.4 Hybrid approaches

Before continuing, another approach is to use filters obtained from either STA/STC or iSTAC and then perform maximum likelihood inference for the parameters of a particular parametric nonlinearity. Such methods can be considered ‘hybrid’ approaches, because they employ fast, moment-based filter estimation in conjunction with maximum likelihood estimation for the nonlinearity. The foremost example of such a method, which we term the excitatory-suppressive pooling (ES-pool) model, was first applied by (Rust et al., 2005) to V1 neurons. In its original formulation, STA/STC analysis was used to derive a set of initial linear filters, which were then sorted into a set of excitatory 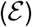 filters (those whose preferred stimulus induced an increase in firing rate) and a set of suppressive 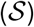 filters (those whose preferred stimulus decreased firing rate). To reduce the high dimensionality of the problem of combining the filter responses, the excitatory and suppressive filter responses were separately pooled via a rooted sum of weighted squares and fed to a fixed nonlinearity *N*(·). This inference process can be summarized as follows:

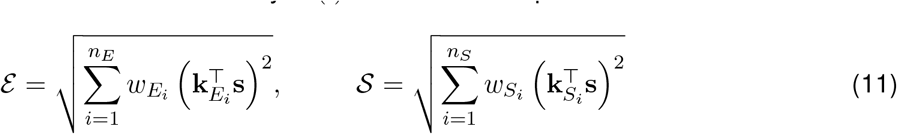

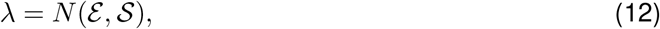

where *n_E_* and *n_S_* are the numbers of excitatory and suppressive filters, respectively, k*_E_i__* and k*_S_i__* are the *i*th filters, and *w_E_i__* and *w_S_i__* are the respective weights for the *i*th filters. λ is interpreted as a Poisson firing rate as in Equation 2. In the original model, a half-wave rectification is applied to the weighted output of the STA filter instead of squaring, but we omit that here for clarity of presentation. The model was then fitted by maximizing the mutual information between the joint excitatory and suppressive signals and the neural response. We modified this initial formulation, however, using iSTAC analysis to derive the initial filters, replacing the static nonlinearity *N* with a feedforward neural network 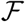, and optimizing the model using gradient descent on the Poisson negative log-likelihood, training both the pooling weights and the neural network parameters. We were able to achieve significantly better performance with these modifications compared to the original model.

## 3 Deep learning models

Deep learning approaches to neural encoding are typically considered distinct from the LNP framework. However, this is not the case. Like the generalized LNP model presented in Figure 1a, a DNN applied to neural encoding accepts an input stimulus and performs a series of affine and nonlinear transformations across its layers before producing a single non-negative value interpreted as a Poisson spike rate. The model can also be constructed to produce multiple outputs, with each value interpreted as a spike rate for a separate neuron in a population (McIntosh et al., 2016). (Note that for clarity’s sake, throughout this article we will use the term DNN to refer to fully-connected feedforward architectures, unless otherwise specified.)

In this way, a DNN is simply a high-dimensional non-linear function approximator (Goodfellow, Bengio, & Courville, 2016). It is this flexibility that enables the unification of these two frameworks, such that a deep network can be seen as the nonlinearity in an LNP architecture, accepting as input the linearly-filtered stimulus (Figure 1b).

Alternatively, an LNP model as a whole can also be seen as a specific type of DNN. Consider a simple LNP model with a single filter **k** ∈ ℝ^*n*^, where *n* is the dimensionality of the input stimulus, and a fixed nonlinearity *f* : ℝ → ℝ_≥0_. Predicting the probability of the spike rate *r* given s, the model can be summarized as

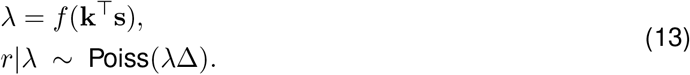

A bias term *b* ∈ ℝ can also optionally be incorporated into the initial affine transformation. Assuming both are trained using gradient ascent on the Poisson log-likelihood, this is exactly equivalent to a 1-layer neural network with parameters *θ* = {**k**, *b*} and activation function *f*. More sophisticated LNP models with multiple filters and fitted nonlinearities, as presented in Equation 8, for example, are also strongly linked to multi-layer DNNs. In this case, the fitted basis functions 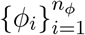 act as the hidden layer(s) of the network, and the 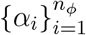 act as the output layer weights.

More precisely, consider a deep network with *L* layers, with each layer *l* structured as

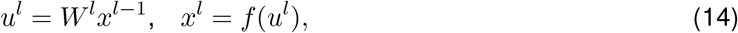

with x^0^ = s. The network is then equivalent to an LNP model with *W*^1^ = *K*^⊤^ and nonlinearity *ϕ* parametrized by *θ* = {*W*^2^, *W*^3^,…, *W*^*L*−1^}, and *α* = *W^L^*:

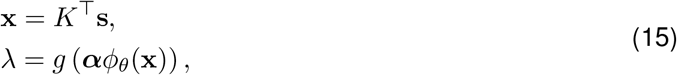

where *g* is a static nonlinearity.

Note, bias terms are omitted here for clarity of presentation, but can easily be added. Interestingly, the canonical formulation does not include a first layer bias term (Williamson et al., 2015), which we found to significantly boost performance when used in DNNs. However, it’s possible that the nature of certain basis functions such as CBFs could implicitly incorporate the computational benefit of a bias through the centers of their Gaussian bumps (see Supplementary section 7.1). In this way, we can see that the basic DNN model is simply a special case of a LNP model with a different parameterization of the nonlinearity. Importantly, the DNN nonlinearity is directly optimized along with the filter weights during training. It is worth noting that it is also possible to reparameterize the input layer of the network as follows:

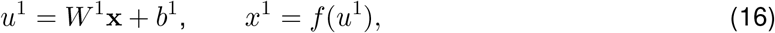

where x = *K*^⊤^ s+z, with z being a bias vector. This effectively adds an extra affine layer before the first layer weights. Interestingly, this formulation has some connection to the *bias transformation* introduced by (Finn, Yu, Zhang, Abbeel, & Levine, 2017), with z acting as the transformation’s parameter vector. Such a parameterization is thought to increase the representational power of the model’s gradient during learning.

If the assumption underlying the LNP framework is correct – namely, that only a few filters are required to cover the relevant stimulus subspace – we would expect that a DNN trained on a neural encoding task should only require several input filters to comprise the first layer weight matrix, regardless of the width of the downstream hidden layers. For example, in a simple cell found in primate V1, STA/STC analysis suggests that approximately 7–8 filters are sufficient to account for the full range of neural responses (Rust et al., 2005). Such bottlenecking of the input is unusual in the construction of DNNs, as typical hidden layer sizes are significantly greater.

Beyond simply reformulating the model, this adaptation has practical effects. Let *W*^1^ *∈* ℝ^*n*×*m*^ be a typical deep network first layer weight matrix, where *n* is the dimensionality of the input stimulus and *m* is the number of output units for that layer. For a DNN constructed with the above assumption, however, if there are *k* < *m* filters, the model will use an *n* × *k* filter matrix followed by a *k* × *m* weight matrix in the next layer. If the second layer weight matrix is normally *m* × *m*, and *k* ≪ *m*, the reduction in the number of parameters is significant. For example, if we have *n* = 500, *k* = 7, and *m* = 100, the model saves 92.9% of the parameters in its first two layers, in theory reducing the risk of overfitting and increasing training speed without losing its ability to capture the correct stimulus response. Additionally, it is possible to simply use iSTAC analysis of the neuron’s spiking response to initialize the input filters (Rust et al., 2005), which further speeds convergence (see Section 5.3). However, results that we present in Section 5 indicate that, contrary to conventional thought, traditional fully-connected DNN architectures are able to achieve a greater correlation with the true spike train through the use of a greater number of input filters. Our subsequent analysis indicates that neurons in the early visual system may have a far more complex response to stimuli than has been previously assumed.

## 4 Experiments

### 4.1 Data

We tested our models on two different neural recording datasets consisting of three different cell types from the retina and primary visual cortex (V1), both obtained *in vivo* from adult macaque monkeys. The first dataset consists of extracellular multi-electrode array recordings from 9 parasol retinal ganglion cells (RGCs), 5 of which were classified as ON cells and 4 as OFF cells. Obtained by (Uzzell & Chichilnisky, 2004), the stimulus consists of a binary full-field flicker generated by a cathode ray tube refreshing at a rate of 120Hz. The second dataset consists of extracellular recordings from 9 simple cells and 9 complex cells from V1. These cells were exposed to a randomized binary bar stimulus aligned with each neuron’s preferred orientation, and the recordings were obtained and the cells classified by (Rust et al., 2005).

For both datasets, we only used cells for which repeat recordings of the same stimuli were obtained as a measure of inherent response reliability, leaving out cells that either lacked such repeat data or for whom reliability was markedly low. We defined a cell as having low reliability if the correlation *r*^2^ between its repeat responses was less than 0.3. Example stimuli from each dataset are viewable in Supplementary Figure 8.

### 4.2 Models

We compared information-theoretic spike-triggered average and covariance (iSTAC), cylindrical basis function (CBF), excitatory-suppressive pooling (ES-pool), and deep neural network (DNN) estimators for all three cell types (RGC, simple, & complex). Due to the relative simplicity of the stimuli, we used only fully-connected neural networks on the retinal data, but for the V1 data we also experimented with 1-D CNNs and recurrent Long Short-Term Memory (LSTM) networks (Hochreiter & Schmidhuber, 1997), designed to process sequential data with long-term time dependencies. Consistent with the findings of (McIntosh et al., 2016), we also found that using larger CNN filters (7-dimensional) than are typically applied in computer vision was more effective.

Each of these models carries with it a different inductive bias governing the functionality of its input filters. In the traditional LNP and fully-connected models, the input filters are applied simultaneously to the concatenated stimuli across all time steps in the relevant stimulus history.

In contrast, in a 1-D CNN, each filter of size *k* < *n*, where *n* is the stimulus dimension, is convolved over the stimuli separately, with the weights shared across time steps. Similarly, the input weights of an RNN are each applied to one stimulus at a time, and shared across time steps. For each model, we ran a hyperparameter search to identify the optimal model width and depth for the DNNs and number of filters and basis functions for the LNP models.

All models were trained using gradient descent on the full Poisson loss function *J*:

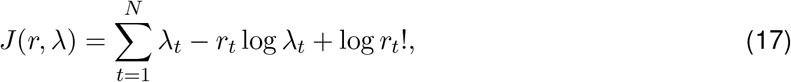

where *N* is the number of time bins, *r_t_* is the true spike count in time bin *t*, and *λ_t_* is the predicted spike rate. The last term, log *r_t_*! is not dependent on the model and is therefore not necessary for training and can be dropped. This is equivalent to minimizing the negative log-likelihood of the neural response under an assumed Poisson distribution. For more model and training details, see Supplementary section 7.2.

### 4.3 Performance Metrics

We experimented with two different performance measures: the single-spike information *I_ss_* and the coefficient of determination *r*^2^. While *I_ss_* is traditionally calculated as *D_KL_* (*p*(s|*spike*)║*p*(s)), where *D_KL_* denotes the KL divergence (Brenner, Strong, Koberle, Bialek, & Steveninck, 2000), (Williamson et al., 2015) showed this is equivalent to

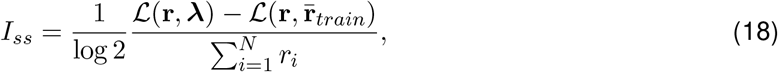

where r is the target spike train. 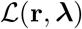 is the model log-likelihood, given by –*J*(*r*, λ) (Equation 17). 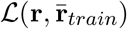 is the log-likelihood obtained by predicting the average spiking rate 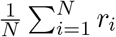, where *N* is the number of bins, in each time bin. The information is then divided by log 2 to convert its units from nats to bits. In this way, maximizing *I_ss_* is equivalent to maximizing the model Poisson likelihood (or, minimizing the negative log-likelihood). It is also worth noting that the quantity 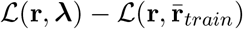 is directly proportional to the *deviance* of the Poisson distribution, a generalization of the use of residuals as a measure of goodness-of-fit to maximum-likelihood models (Nelder & Wedderburn, 1972). Using the coefficient of determination, or *r*^2^, simply measures the proportion of the variance in the true spike response that is explained by a given model. In addition to being a simple and commonly used metric, it is numerically unrelated to the single spike information. The use of repeat data, in which the same stimulus is replayed over multiple trials to the same cell, allows us to compare the correlation of a cell’s responses with itself to those obtained by our models. We found that *I_ss_* and *r*^2^ were generally closely correlated, and generally default to presenting results in terms of *r*^2^, both for the ability to upper bound them via the use of repeat data, as well as due to its greater use in the wider literature.

## 5 Results

### 5.1 Retinal Ganglion Cells

We trained traditional LNP nonlinearities as well as DNN nonlinearities on the RGC data. Due to the simpler nature of both the full-field flicker stimuli and the retinal cellular responses themselves, we only used feedforward fully-connected (FC) networks. We found that a simple 2-layer network was enough to outperform the traditional LNP nonlinearities. These results were in accordance with the prediction that performance saturates with a relatively low number of filters (Figure 2). Average performance of the best model for each nonlinearity can be seen in Figure 4a. Examining the 1-D filters learned from the same example ON cell whose performance is plotted in Figure 2b, we can see that the network learns similar filters to those obtained through STA/STC analysis, with the differences accumulating from left to right. In particular, the primary iSTAC filter, which accounts for the most information in the cellular response, is found nearly exactly by each of the other nonlinearities. One notable feature of the DNN filters is their relative broadness and simplicity in comparison to the other models. While the other nonlinearities learn filters that have multiple closely concentrated peaks, the corresponding DNN often has a single broad area of activation in the same location. The effect is that the DNN is less discriminating in responding to stimuli, resulting in a broader nonlinearity (see Supplementary Figure 9a). It appears that the neural network nonlinearity is more sensitive to a wider range of the stimulus space, with the traditional LNP nonlinearities being more selective.

**Figure 2:**
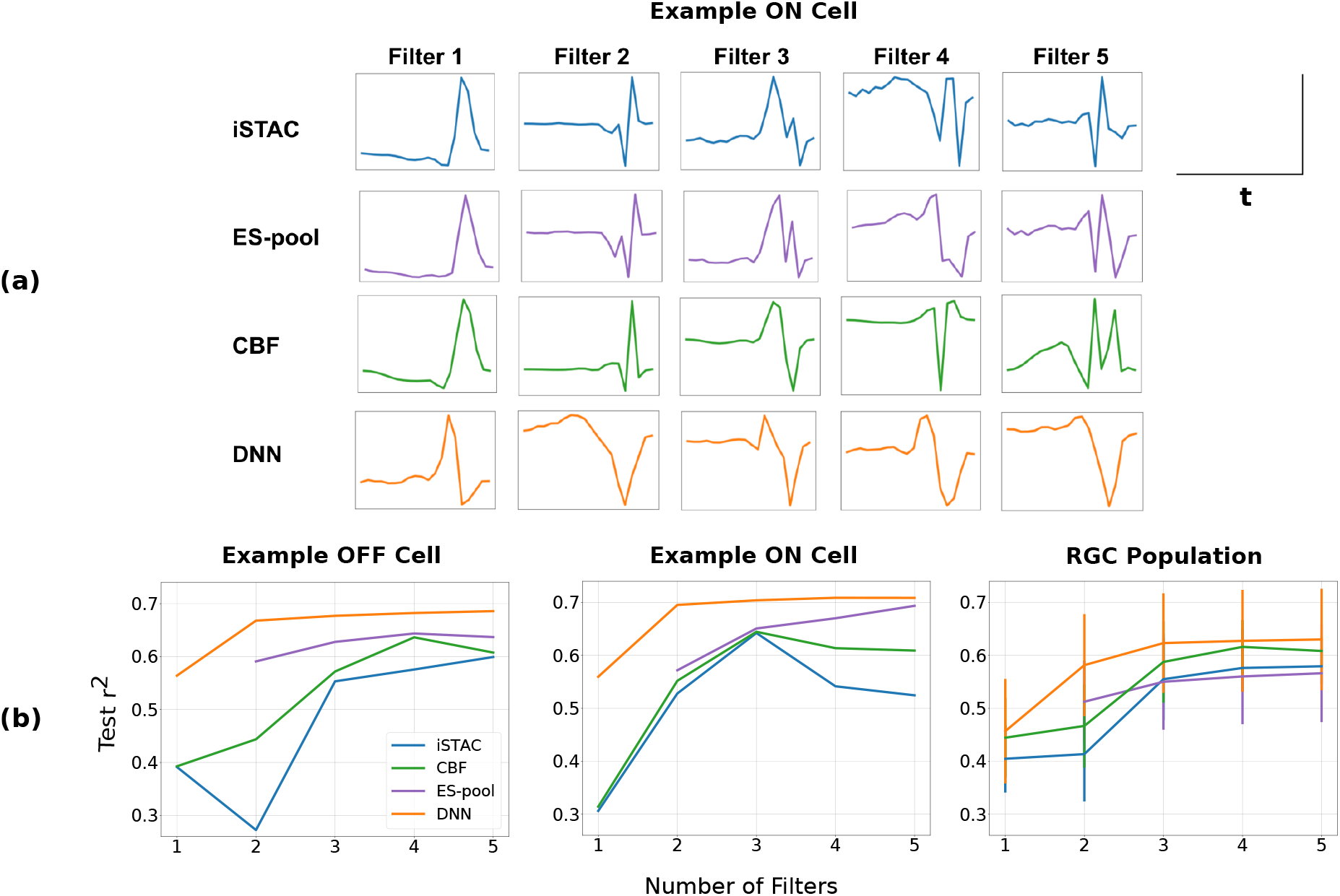
RGC cell results. (a) The 1D temporal filters learned by each estimator, ordered by alignment with the sorted iSTAC filters. In contrast to the other models, the DNN filters are more generalized, usually learning one broad peak where the other model filters learn a cluster of narrow spikes. (b) Model performance for an example ON cell, example OFF cell, and the average across the population. The example ON cell is the same as in part (a). Note that the Excitatory-Suppressive Pooling (ES-pool) model results begin with two filters, as the model fundamentally relies on two streams of processing. The DNNs display higher performance than the other methods, though for all nonlinearities performance saturates after approximately five filters.

### 5.2 V1 Cells

The differences between the DNN models and the traditional LNP nonlinearities are more apparent in the V1 results. While for equal numbers of filters (Figure 3b), the fully-connected DNN performs on par with the other LNP nonlinearities, it’s able to continue improving beyond the point at which performance saturates for the other models (Figure 5). This difference becomes more substantial as the complexity of the filters grows (Figure 4). It is likely that the relative non-specificity of the DNN filters contributes to this comparative lack of a performance ceiling (note the relative lack of smoothness in the DNN filters in Figure 3a). Although absolute test performance is low in the V1 cells, it is still comparable to that achieved by the retinal models relative to the performance ceiling established by the repeat data. While difficult to discern in the complex cell nonlinearities, the same relative broadness in the plotted nonlinearity in comparison to the traditional models can be seen in Supplementary Figure 9b,c.

**Figure 3:**
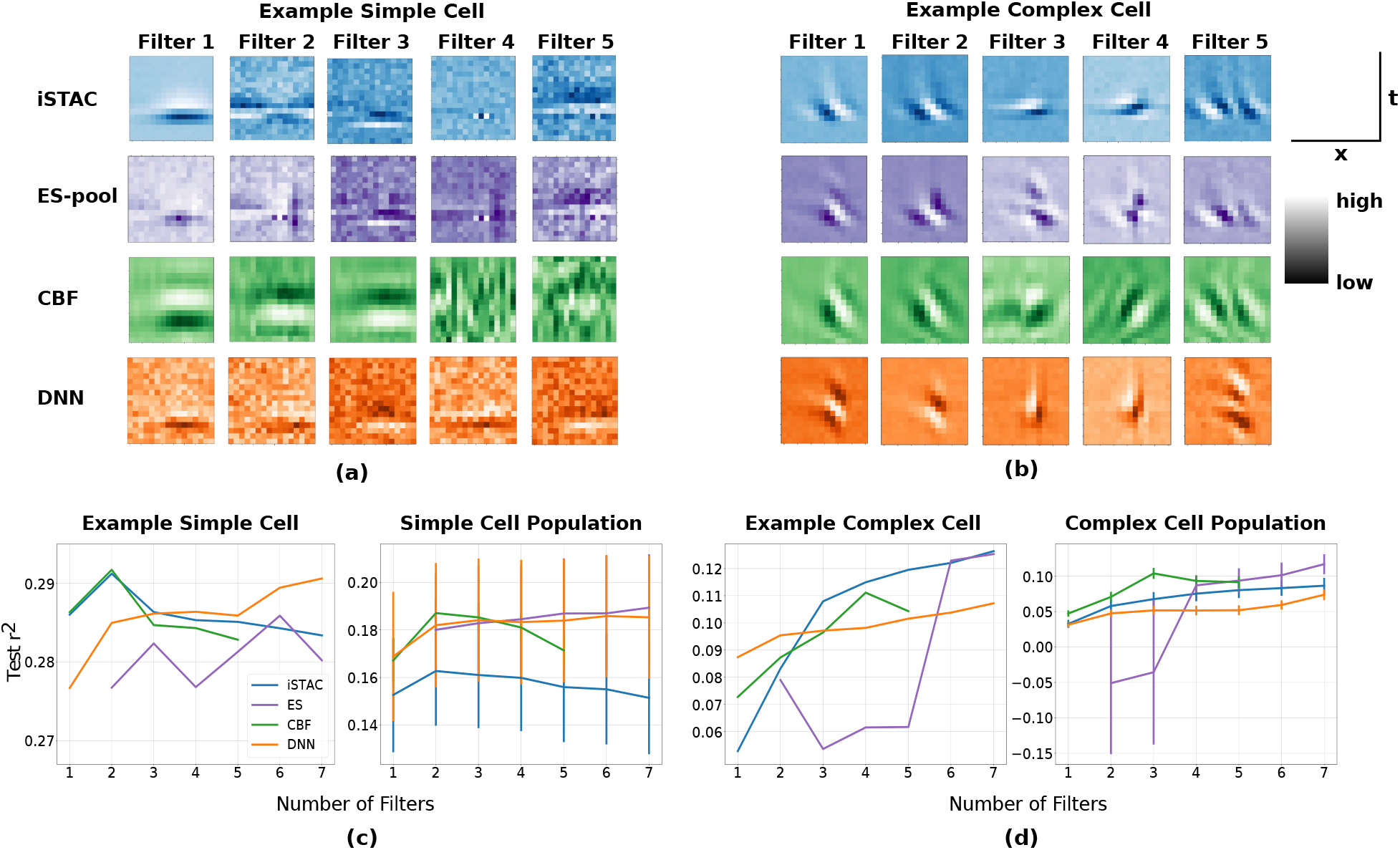
V1 cell results. (a) The spatiotemporal filters learned by each estimator, ordered by alignment with the sorted iSTAC filters. The models learn similar filters, although in general the DNN filters exhibit less smoothness. (b) Average model performance across the population and on an example cell for each cell type as a function of the number of input filters in the model. Note that the Excitatory-Suppressive pooling (ES-pool) model results begin with two filters, as the model fundamentally relies on two streams of processing. The high variability is a result of high variability in the reliability of the cellular response. Most models typically reach peak performance at seven filters or fewer, but DNNs continue to improve beyond that number (see Figure 5).

**Figure 4:**
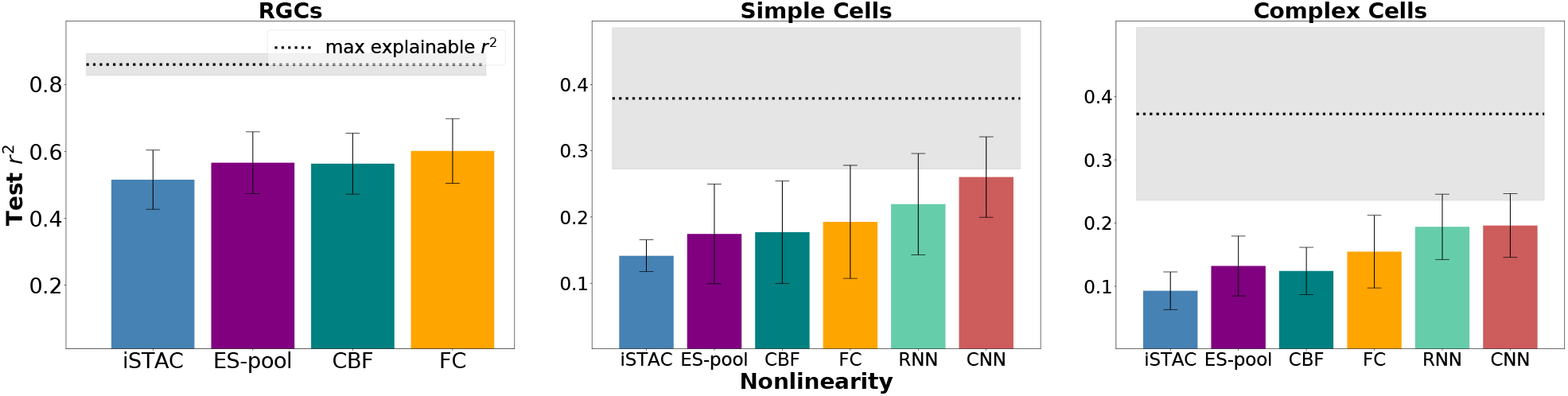
Test *r*^2^ means and standard deviations for the best model of each type trained on (a) RGCs, (b) simple cells, and (c) complex cells. The average *max explainable r*^2^ is the *r*^2^ between two halves of the repeat data, averaged across different splits of the data and across cells. This acts as a measure of the average reliability of the population’s response, effectively upper-bounding possible model performance. The shaded region is the standard deviation of these value across cells. The neural network models display an increasing advantage over the traditional LNP nonlinearities as the complexity of the cellular responses and input stimuli increases.

**Figure 5:**
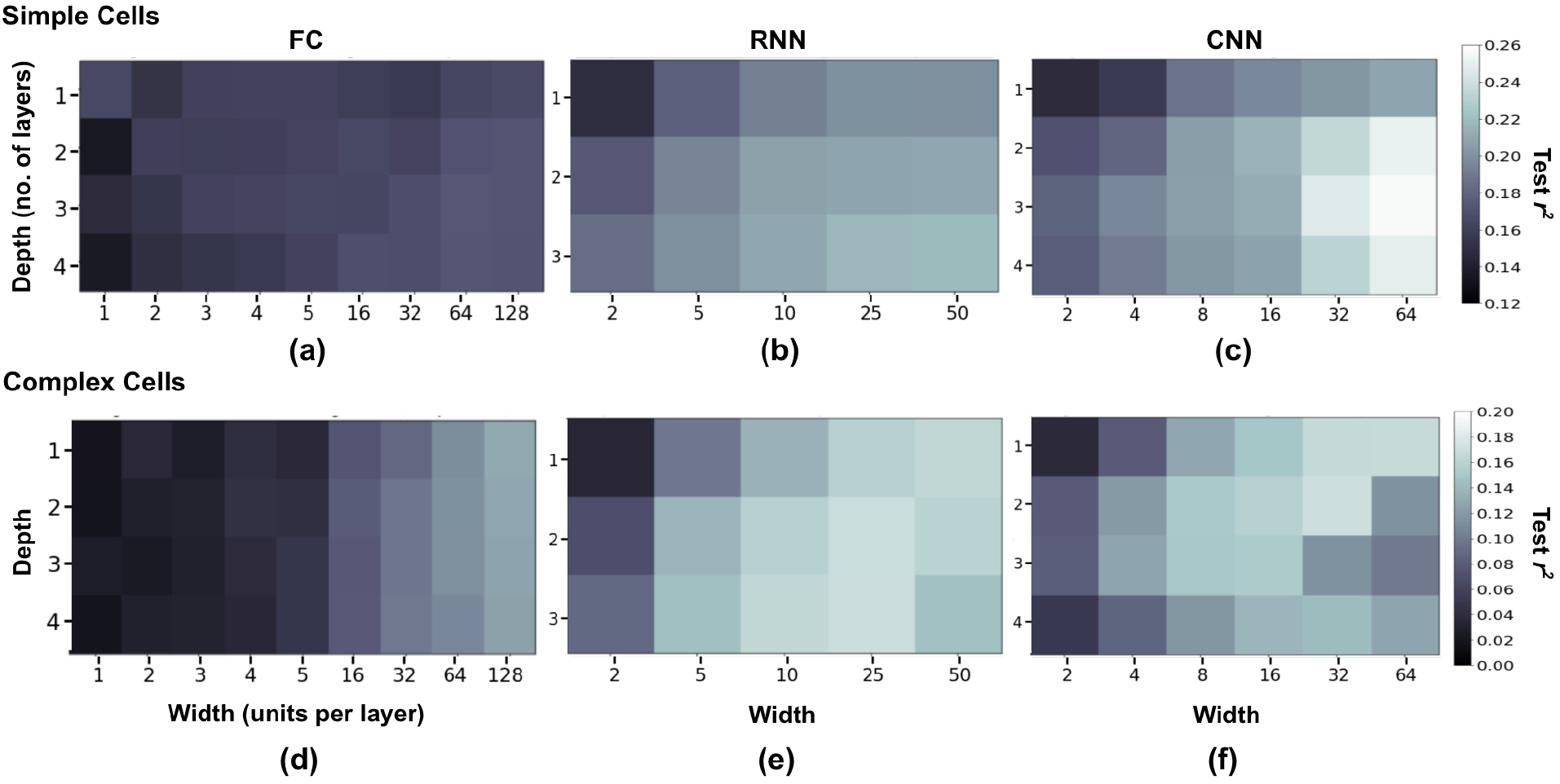
Average held-out test performance as a function of the depth and layer width of neural network models for simple cells (a-c) and complex cells (d-f). Each grid square represents the average final test *r*^2^ across each cell in the population. Each architecture had a constant number of units per layer with the exception of the output layer. For fully-connected (FC) models, the results show that performance improves consistently as a function of model width, with depth having a more moderate effect. In recurrent (RNN) and convolutional (CNN) models, there is a more complex relationship between architecture and performance.

The fully-connected network performance continues to improve beyond 7-8 filters as shown in Figure 5. This challenges the notion that only a few filters are required to fully characterize the neural response. However, one possible reason that these larger models outperform the others is not that the cellular response is indeed sensitive to a high-dimensional subspace, but that the models require the greater number of parameters in the larger model to adequately learn a smaller subspace. To investigate whether the latter is true, or if a large number of filters are actually required to characterize the neural response, we performed principle components analysis (PCA) on the input filters of the best fully-connected and convolutional models trained on the simple and complex cell populations. We then reconstructed the full filter matrix one principle component (PC) at a time, evaluating the model’s test performance after each component was added. As a form of normalization to allow comparability across cells and models, we divided the *r*^2^ performance of each model by their peak performance as PCs were added to the reconstruction. If the relevant stimulus subspace is in fact low-dimensional, then the performance of a large, trained network should not suffer if it filters the input with only the first few PCs of the input weights. If a higher dimensional stimulus subspace affects the neural response, then accordingly a higher dimensional filter space must be learned to adequately filter it, a greater number of PCs would be required to reach optimal performance.

The results are summarized in Figure 6. We can see that for many of the simple cells, while high performance is obtained very quickly, it peaks at ~10 PCs and declines or levels off quickly as PCs continue to be added in the fully-connected network. This indicates that for these cells the relevant subspace likely has lower dimensionality than the complete number of filters, but that it still has higher dimensionality than the LNP framework typically assumes. Indeed, there are several cells for which performance continues to increase as components are added (Figure 6a). On average across the population, performance is relatively flat and even declines slightly as PCs are added (Figure 6c). Especially strong performance on those simple cells whose performance does improve as PCs are added is then likely why average FC performance is slightly above the traditional nonlinearities for simple cells (Figure 4b), as for the low-dimensional cells it underperforms. This is an important result, as it shows that there is considerable variability even within cell types as to the nature of the neural response. It would be useful in the future to test this trait in a larger population of cells. Therefore while the low-dimensional hypothesis seems to hold in whole or in part for many of these cells, for many it appears to fail.

**Figure 6:**
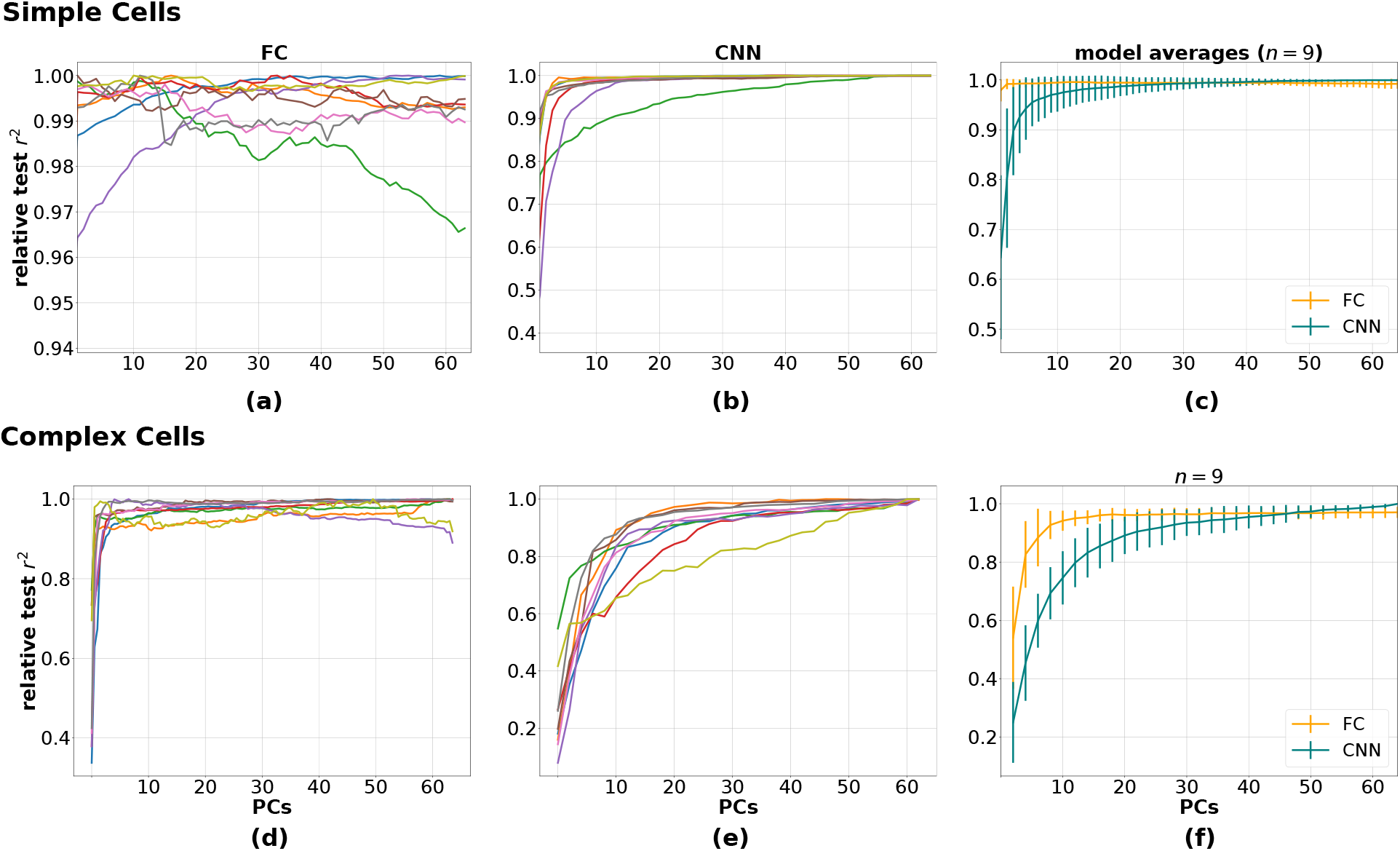
Normalized test performance for fully-connected models (a,d), convolutional models (b,e), and on average across the cell populations (c,f) as a function of the number of principle components (PCs) used to reconstruct the input filters. Each trace in parts a,b and d,e represents a separate cell. The top row is simple cell results and the bottom is complex cells. To standardize across cells, we divided each curve by the maximum *r*^2^ value obtained as PCs were added. If the low-dimensional encoding hypothesis is correct, we would expect to see performance peak at <10 PCs. For most cells, this is not the case.

In contrast to the simple cells, nearly every fully-connected complex cell model improves performance as components are added. This indicates that complex cells likely do respond to a much higher-dimensional stimulus subspace, and could also partly explain the superior performance of FCs over traditional nonlinearities (Figure 4c). Moreover, across both cell types, CNNs improve performance as PCs are added (Figure 6b,c). This is likely because each 7-dimensional kernel learns a specific feature of the simple 16-dimensional stimulus space, and so a greater number of filters are required to cover it. It is also likely that for higher dimensional natural stimuli that performance would not begin to saturate as seen here.

### 5.3 Model Efficiency

We also tested several variants of the bottlenecked DNN framework discussed in Section 3. In addition to the ES-pool model, we trained fully-connected DNNs with their initial filters determined through iSTAC analysis. We tested two conditions within this strategy: one in which these filters were kept frozen during training, and one in which the filters were allowed to train. The second condition produced far superior performance, and in general initialization using iSTAC filters nearly halved model training time in comparison to random filter initialization in the standard DNN models (Figure 7). In addition, we not only measured the single-spike information (*I_ss_*) of each model, but on average how much information was accounted for per model parameter (*I_ss_*/parameter), shown in Supplementary Figure 10a,b. Interpreted as a measure of model efficiency, we can see that all of the low-filter DNN variants outperform the standard DNN by this measure, with the ES-pool model being especially notable.

**Figure 7:**
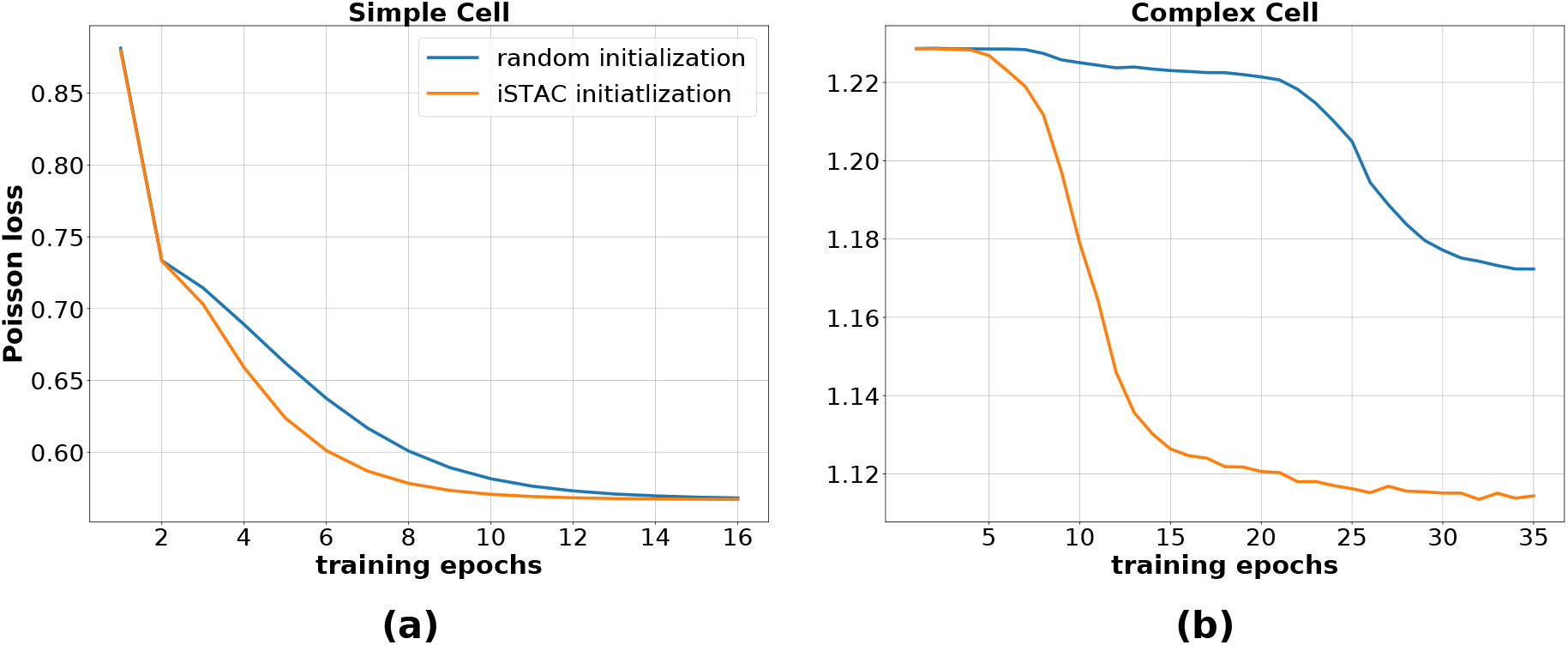
When possible, initializing filters with iSTAC analysis can have a significant benefit to minimizing both loss and convergence time. These results are from fully-connected models. (a) In this simple cell, such initialization does not make a difference in the final performance, but it does nearly cut convergence time in half. (b) Not only is convergence time reduced in this complex cell, but the overall performance is significantly improved via iSTAC initialization. The benefits of iSTAC initialization are unsurprising, as it the standard procedure for more traditional LNP estimators.

## 6 Discussion

Collectively, these results demonstrate both the fundamental compatibility of the DNN and LNP approaches, as well as show that as nonlinear function approximators, DNNs have a greater capacity to exploit all of the relevant information in the stimulus space compared to more traditional nonlinearities. While their feedforward architectures may fit within the LNP framework, the specialized optimizers used to perform gradient descent in deep models also confer an advantage. Through the CBF nonlinearity, these results also connect deep networks to information-theoretic estimators like MID. This compatibility also allows for easier comparison between model components. Examining the filters and resulting nonlinearities above shows that DNNs learn a more diffuse representation of the stimulus subspace and as a result are able to capture a broader range of the neural response. It should be noted that these results do not completely overturn the understanding that only a small number of filters are necessary to characterize the relevant stimulus subspace for neurons in the early visual system. It is clear that the vast majority of performance gains, as well as the bulk of the relevant information, is obtained in the first few filters, even for cells that seem to require a high number of filters for optimal performance. However, they do indicate that DNNs are better able to take advantage of residual information in the stimulus. Similar gains were not observed in the traditional LNP nonlinearities when the number of filters was increased beyond its usual limit, although it was not computationally tractable to push the number of filters to the same level as the DNN models. This computational efficiency is another factor in favor of their use. It is also clear that the different ways in which RNNs and CNNs process the input produces significant advantages over other models, although the reasons for this have no yet been fully explored.

The fact that these effects are observed even for simple binary stimuli implies not only that neurons are sensitive to a richer subspace of the input stimulus than previously thought, but also that these discrepancies are likely even more significant for natural stimuli. Investigating the underlying causes of these differences is a promising avenue for future work and a greater understanding of neural encoding.

## 7 Supplementary Information

### 7.1 First Layer Biases

It is possible that, while the traditional LNP formulation does not include a first ‘layer’ bias to accompany the linear transformation of the input stimulus, certain basis functions may play a similar role. For example, a ‘first-order’ CBF is defined as a Gaussian bump aligned with a particular direction of the feature space, centered at *μ* with characteristic width *σ*:

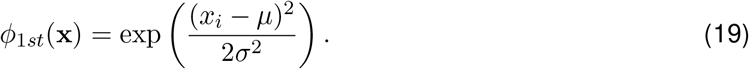

Viewed alternately as acting on the shifted stimulus projection *x_i_*–*μ*, we can see that the CBF effectively applies a uniform first layer bias term. Defining a vector of such centers *μ* ∈ ℝ^*k*^, where *k* is the number of filters, with *μ_i_* used to offset *x_i_*, would be equivalent the unconstrained first layer biases used by DNNs.

### 7.2 Model and Training Details

All neural network models used the softplus nonlinearity at each layer and were trained using the Adam optimizer (Kingma & Ba, 2014). The softplus nonlinearity was chosen for numerical stability in conjunction with the Poisson loss function. Dropout (Srivastava, Hinton, Krizhevsky, Sutskever, & Salakhutdinov, 2014) was added after hidden layers of sufficient size to prevent overfitting. Simple cell model architecture details can be found in Table 1.

**Table 1:**
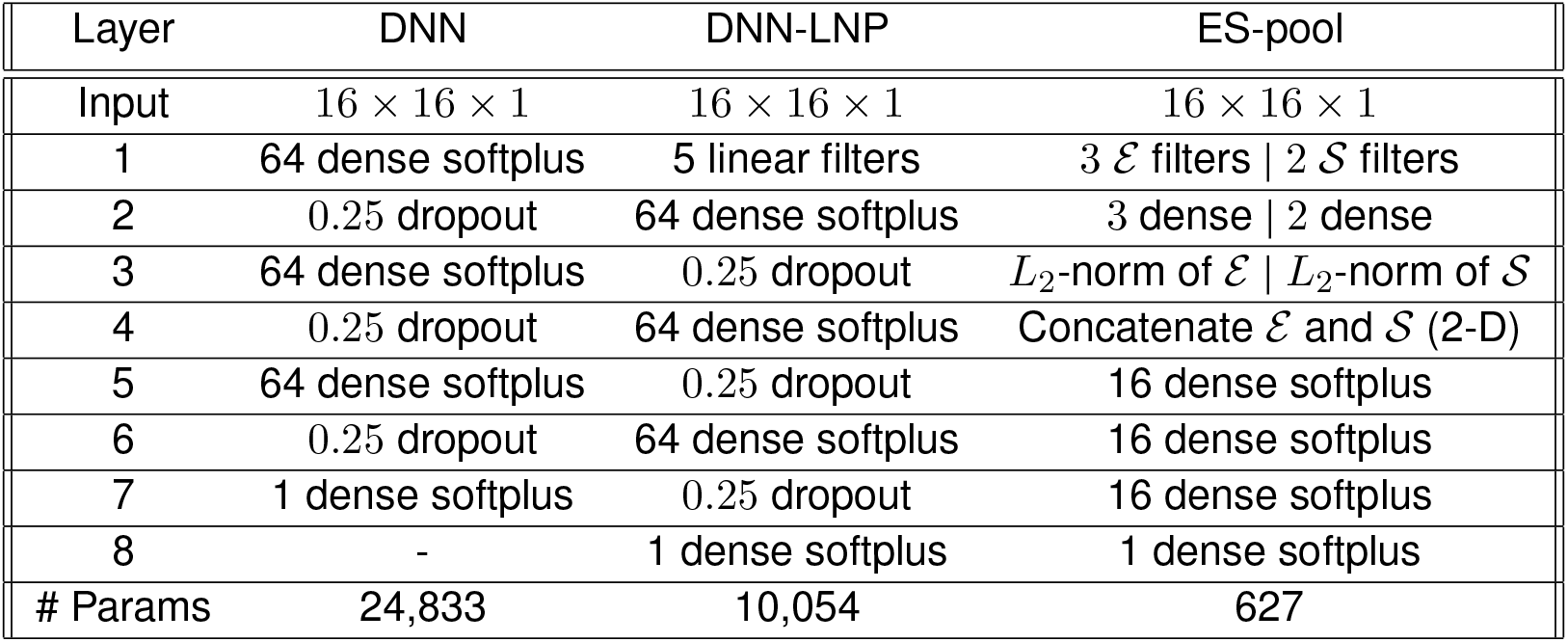
Several simple cell model architectures. Input was flattened to 256 × 1. ‘# Params’ refers to the number of trainable model parameters. A ‘dense’ layer is a fully-connected layer. DNN-LNP refers to a model with an extra linear layer for dimensionality reduction as described in Section 3.

### 7.3 Figures

**Figure 8:**
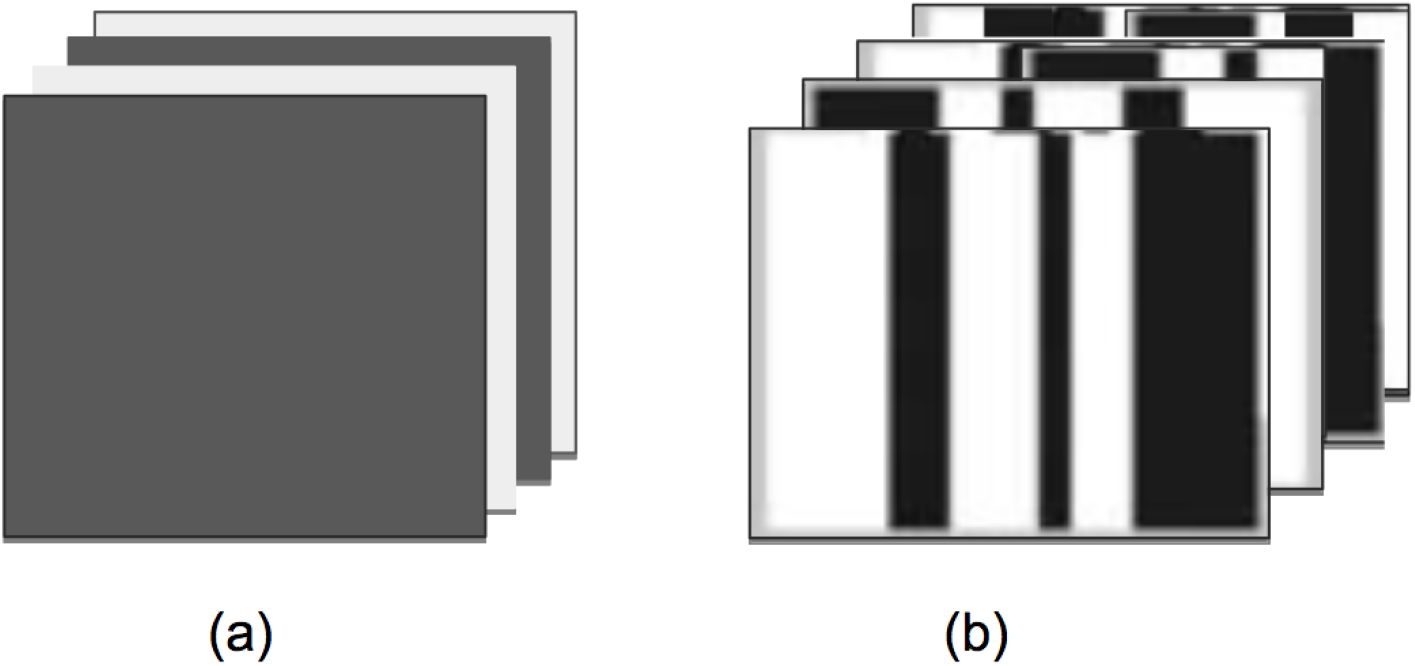
Sample stimuli. (a) The retinal data stimuli simply consisted of binary full visual field flickers. (b) The V1 stimuli were only slightly more complex, consisting of binary bars aligned with the target neuron’s preferred direction of orientation. The fact that the most successful V1 models required a high number of input filters even for such simple stimuli implies that the fundamental dimensionality of simple and complex cell responses is higher than previously thought.

**Figure 9:**
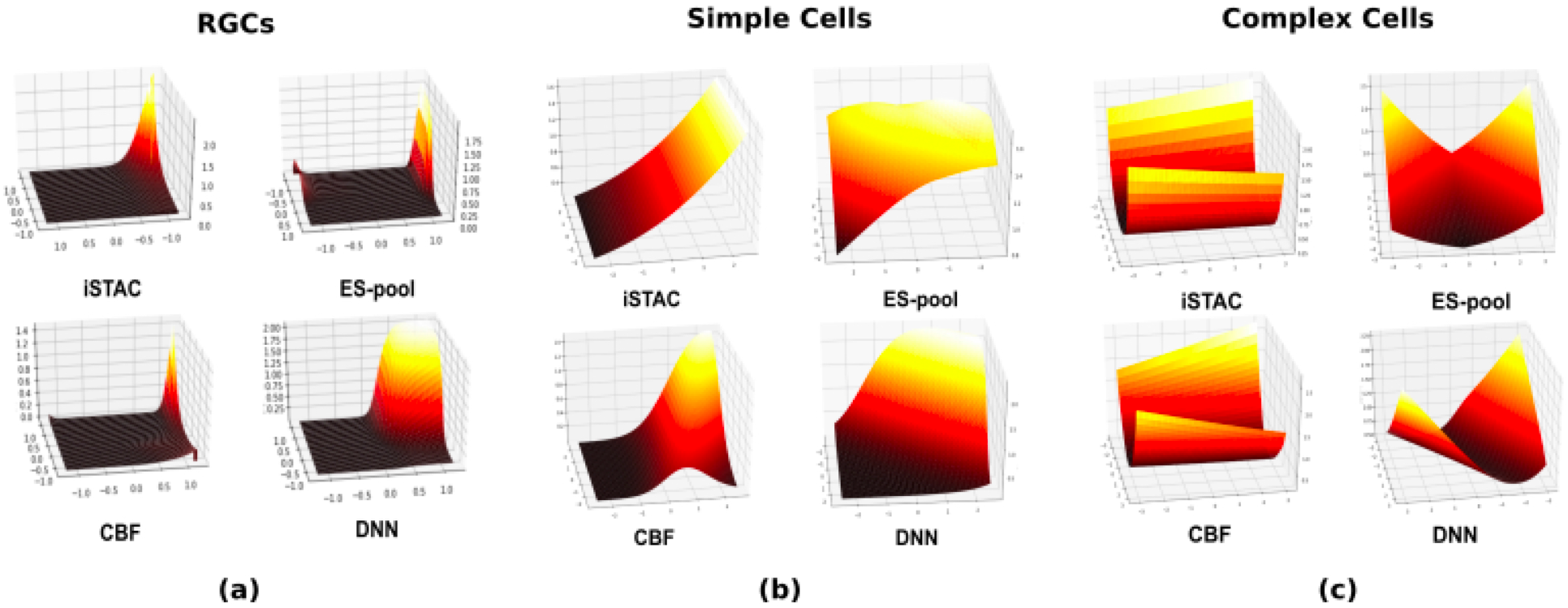
Nonlinearities from two-filter models. The two filters were orthogonalized and used to create a basis from which a 2D grid of stimuli was generated. The model firing rates in responses to these stimuli form the nonlinearity. It should be noted that for all nonlinearities other than those trained on RGCs, two-filter model performance was very weak. (a) Models trained on RGCs. These plots are the most interpretable, clearly demonstrating the relative broadness of the DNN nonlinearity relative to the other models. This is likely a direct consequence of the relative broadness of the DNN filter contours in Figure 2a. (b) Simple and (c) complex cell nonlinearities. These results are more difficult to interpret, likely because two-filter models trained on these data did not achieve high performance.

**Figure 10.**
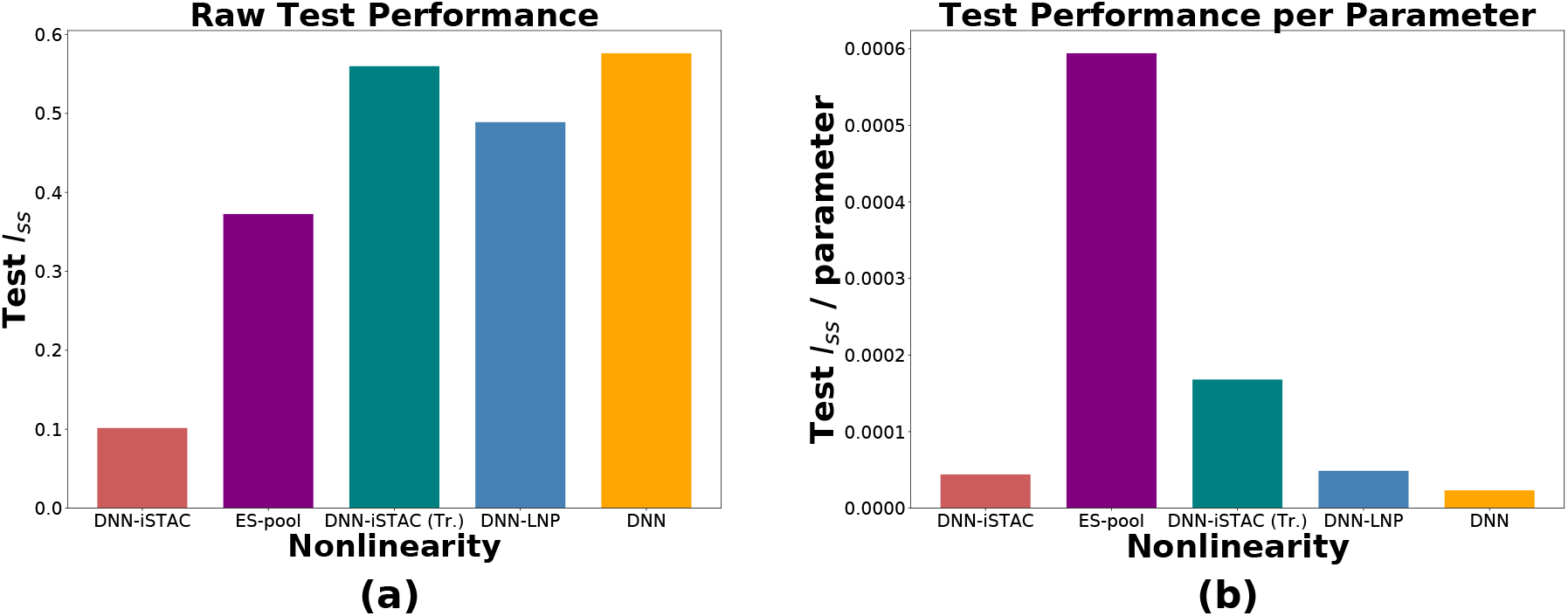
Measures of model efficiency: (a) test single spike information (*I_ss_*) and (b) test single spike information per parameter (*I_ss_*/parameter) for simple cell DNN model variants. Here, DNN-iSTAC denotes models whose input filters were initialized with iSTAC filters, but were frozen during training, while DNN-iSTAC (Tr.) denotes models whose filters with initialized with iSTAC but were allowed to train. DNN-LNP denotes models with an additional linear layer for dimensionality reduction as described in Section 3. We can see that these and the ES-pool models achieve significantly greater parameter efficiency.

